# Efficient Methods for Target Gene Manipulation in Haematopoietic Stem Cell Derived Human Neutrophils

**DOI:** 10.1101/2023.06.17.545406

**Authors:** Ng Anthony Yong Kheng Cordero, Eleonore Fox, Jack Brelstaff, Mattia Frontini, Charlotte Summers

**Affiliations:** *Victor Phillip Dahdaleh Heart and Lung Research Institute, University of Cambridge, Papworth Road, Cambridge, CB2 0BB, United Kingdom.; Department of Haematology, NHS Blood and Transplant Centre, University of Cambridge, Long Road, Cambridge, CB2 0PT, United Kingdom.

## Abstract

Neutrophils are the most abundant leukocyte in humans and the principal effectors of the innate immue response. Genetic modification of human neutrophils is challenging due to their short lifespan and tendency to activate in response to even minor perturbation. However, genetic manipulation of haematopoietic progenitor cells and subsequent directed differentation into neutrophils represents a potential avenue to study the contributions of individual genes and pathways to human neutrophil function. Here we present a method of directed granulocytic CD34^+^ progenitor differentiation into neutrophils capable of key functions such as priming and neutrophil extracellular trap (NET) formation. We further show that differentiating progenitors can be efficiently and stably modified by lentiviral gene delivery and Cas9-gRNP nucleofection to produce potent and activation-free gene knockdown in mature neutrophils, thereby providing new tools for understanding the contribution of neutrophils to health and disease. Using this model we have shown that, contrary to previous reports, CD11b is not required for phagocytosis of serum-opsonised bacterial particles.

## 1 Introduction

Neutrophils are the principal effectors of the innate immune response and the most abundant leukocyte in humans, responsible for defending the host against invading pathogens ^1^. Following their maturation from myeloid precursors within the bone marrow, neutrophils enter the peripheral circulation and distribute between the circulating and marginated (intravascular) pools. Neutrophils are short lived cells, with a circulating half-life of six to eight hours ^2^ in humans. In culture, neutrophils undergo spontaneous apoptosis within 24-48 hours. Due to their short lifespan and propensity to become activated when handled *in vitro*, mature neutrophils have proved challenging to transfect or genetically modify, limiting the study of the contribution of specific genes and pathways to neutrophil functions in health and disease. This significant technical challenge has limited our understanding of the molecular mechanisms of neutrophils in health, infection, and inflammatory diseases such as acute respiratory distress syndrome.

Current models used to study neutrophil function include cell lines such as dimethylsulfoxide (DMSO) differentiated HL-60 cells, and animal models such as mice and zebra fish. Differentiated HL-60 cells exhibit significant differences to human neutrophils at both transcriptional ^3^ and functional levels ^4, 5^. Genetic manipulation of animal models is costly and labour-intensive, and does little to overcome the problem of the cross-species differences in the form and function of neutrophils. Robust methods for the genetic manipulation of primary human neutrophils are currently lacking.

Recent advances in genome editing using viral vectors and clustered regularly interspaced short palindromic repeat-Cas9 (CRISPR-Cas9) technology provide a promising avenue for the precise genome editing of CD34^+^ haematopoietic stem and progenitor cells (HSPC). Current approaches for efficient delivery of genetic material to HSPC include the use of viral vectors such as adeno-associated virus six ^6^ or lentiviruses ^7^, and electroporation or nucleofection ^8^. Although these methods result in priming and cell death if attempted in mature circulating neutrophils, they are well tolerated by progenitors cells. Both lentiviral vectors and CRISPR-Cas9 technology rapidly effect stable genetic alterations in the target cells, which persist long after the active editing machinery is degraded, posing little risk of editing-related cellular activation of sensitive cells such as neutrophils.

Here, we describe the rapid and efficient generation of a highly pure population of genetically edited, mature human neutrophils, achieved by *ex vivo* expansion and differentiation of isolated human CD34^+^ HSPC into segmented cells with mature neutrophil functional capacities, and the stable and efficient lentiviral transduction of differentiating progeny. Further, we demonstrate that CRISPR-Cas9 gene editing of differentiating progenitors can be used to facilitate the study of molecular basis of human neutrophil function.

## 2 Results

### 2.1 Differentiated CD34^+^ cells recapitulate multiple mature neutrophil functions

We first sought to establish a robust model for differentiation of CD34^+^ HSPC into human neutrophils. CD34^+^ HSPC isolated from human apheresis cones by magnetic selection were expanded in serum-free medium supplemented with recombinant human interleukin 3 (IL-3), interleukin 6 (IL-6), thrombopoietin (TPO), stem cell factor (SCF), and human fms-like tyrosine kinase 3 ligand (Flt3L) for six days, followed by directed differentiation in 10 ng/mL of recombinant human granulocyte colony stimulating factor (G-CSF) for up to 16 further days. Representative cytospins of unenriched cultures at different stages of G-CSF directed differentiation are shown in Fig. 1A.

**Fig. 1.**
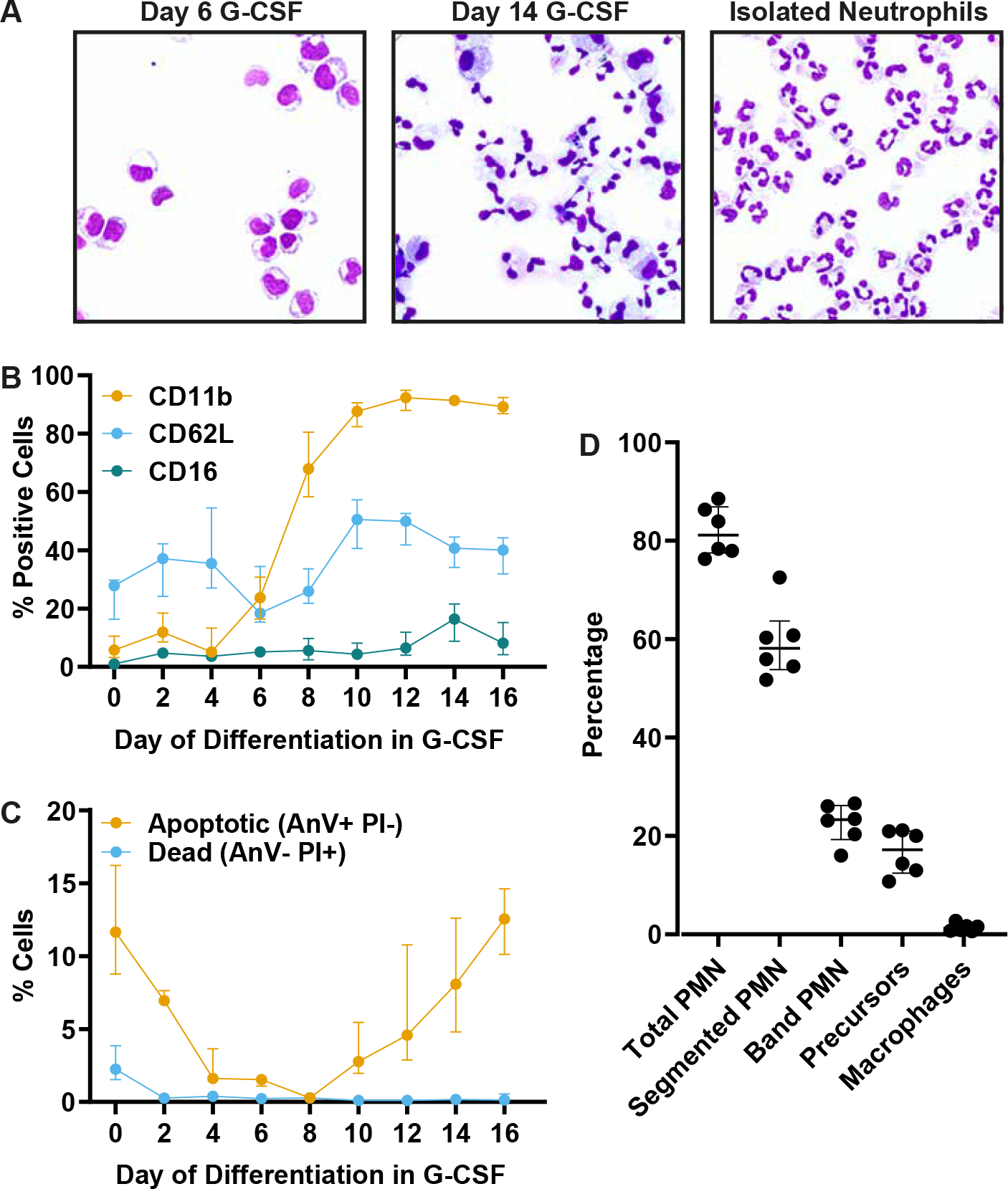
**A** Representative images of differentiating CD34^+^ cells at the progenitor stage (day 6) and as segmented cells (day 14) during directed differentiation with G-CSF. The rightmost image shows isolated mature human neutrophils. Cells were deposited onto glass slides, fixed, and stained with a modified Romaowsky stain. **B, C** Expression of neutrophil surface markers (B) and apoptotic cells (C) during directed differentiation with G-CSF. Cells were stained for 30 minutes on ice and analysed on a BD LSR Fortessa flow cytometer. **D**Percentage of cells in culture after directed differentiation. Total PMN is the sum of band and segmented neutrophils. Cytospins were imaged and counted manually using ImageJ software. At least 1000 cells from 10 different images were counted to determine percentages. Data for B-D are shown as individual data from six distinct donors (dots), with horizontal bars indicating the median of each group with the interquartile range depicted by error bars.

During directed differentiation, segmented neutrophils were first observed from day 10, and were most abundant at day 14. Flow cytometric analysis of myeloid markers (Fig. 1B) showed an increase in surface CD11b expression from day 6 onwards, peaking by day 8 and remaining elevated to day 16. CD62L showed biphasic expression with an initial peak at day 4, followed by a sustained upregulation from day 10 onwards. CD16, which marks mature circulating neutrophils, peaked at day 14. Apoptotic cells (Fig. 1C) were present in decreasing levels from day 0-4, and began to increase from day 10 onward, peaking at day 16, while necrotic cells were infrequent during differentiation. Numbers of recoverable cells in culture decreased strikingly after day 16, in keeping with the apoptosis rates observed. Unenriched cultures at day 14 contained 81.9% neutrophils (59.2% segmented neutrophils, 22.6% band cells), 16.7% myeloid precursors, and 1.4% macrophages or other myeloid cells (Fig. 1D). By day 14, differentiated CD34^+^ cells showed segmented nuclear morphology similar to isolated circulating neutrophils (Fig. 1A). Based on these findings, we selected 14 days of directed differentiation in G-CSF as the optimal duration prior to phenotypic analysis of differentiated CD34^+^ cells.

We next assessed the function of differentiated CD34^+^ cells in comparison to freshly isolated human neutrophils. Pre-incubation of differentiated CD34^+^ cells with platelet activating factor (PAF) followed by the formyl peptide receptor agonist formyl-Met-Leu-Phe (fMLP) resulted in an oxidative burst of similar kinetics, but lower magnitude, to isolated healthy neutrophils (Fig. 2A). To assess phagocytic capacity, isolated human neutrophils or differentiated CD34^+^ cells were incubated with serum-opsonised fluorescent *Staphylococcus aureus* or *Escherichia coli* bioparticles in the presence of the dihydrorhodamine, a fluorescent probe for reactive oxygen species. Differentiated CD34^+^ cells showed phagocytic responses of a similar magnitude to isolated neutrophils. In response to *E coli* bioparticles, differentiated CD34^+^ cells had a reduced oxidative response (Fig. 2B-D).

**Fig. 2.**
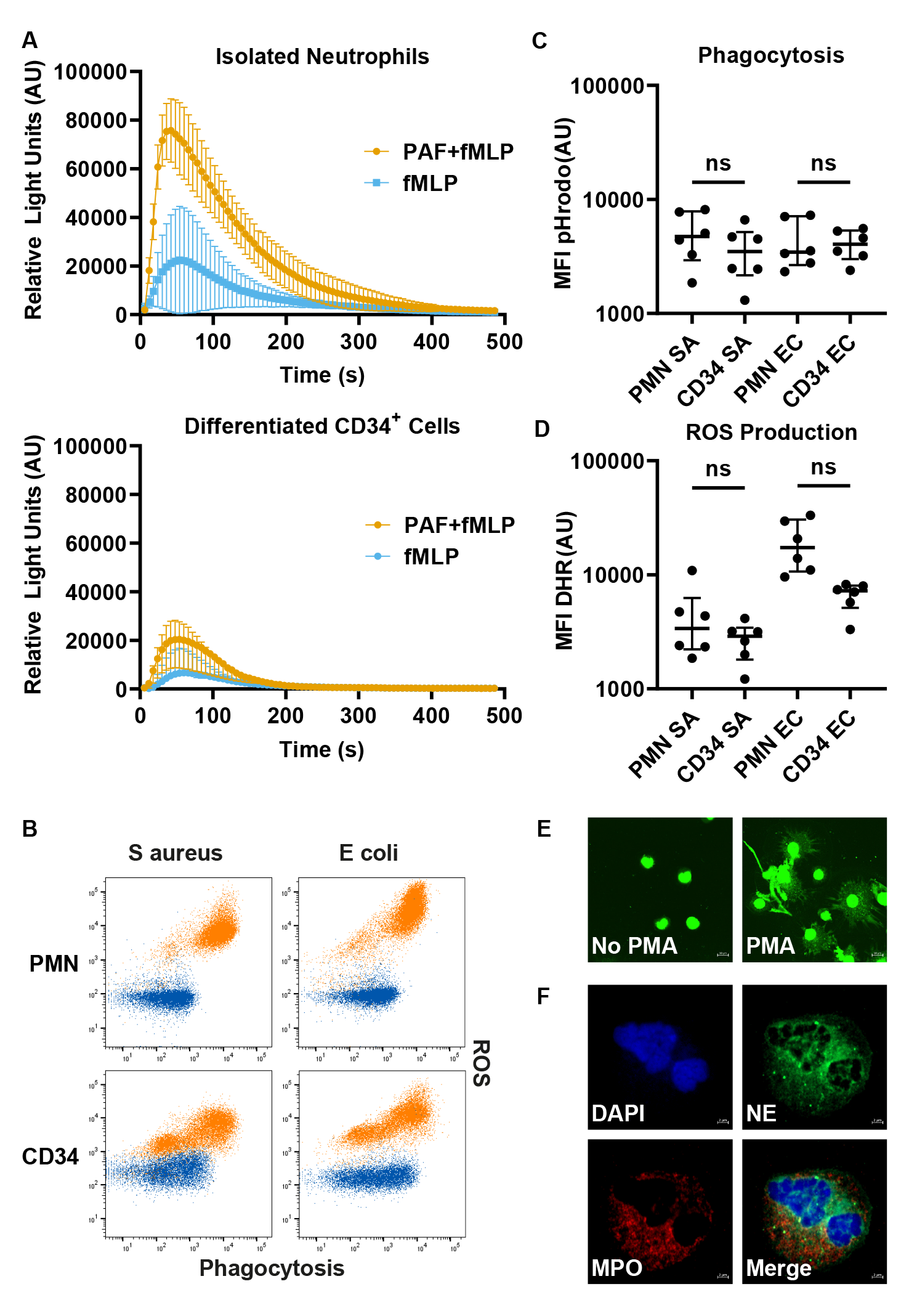
**A** fMLP-stimulated respiratory burst measured by chemiluminescence in differentiated CD34^+^ cells (left) and neutrophils (PMN, right). 4.5 *×* 10^5^ cells were primed with PAF (orange) or mock-primed vehicle (blue) for 5 minutes. Primed or mock-primed cells were incubated with luminol and horseradish peroxidase (HRP) and stimulated with fMLP. Luminescence measured at 5 second intervals after stimulation. Data are in relative light units (RLU) presented as mean and standard deviation of three distinct donors. **B** 4.5 *×* 10^5^ cells were incubated for 30 minutes with 25 *µ*g/mL of serum opsonised *Staphylococcus aureus* (SA) or *Escherichia coli* (EC) bioparticles in the presence of 3 *µ*M dihydrorhodamine and analysed on an LSR Fortessa flow cytometer. Experiments incubated at 37°C are shown in orange, and identical cold controls at 4°C are shown in blue. Events shown are of all live cells (Fig. S1B). **C-D** Differentiated CD34^+^ cells and human neutrophil phagocytosis (C) and oxidative responses (D) from experiments depicted in B. Data from six distinct donors are shown as dots, with horizontal bars indicating the median of each group with the interquartile range depicted by error bars. Data were analysed with the Kruskal-Wallis test between PMN and differentiated CD34^+^ cells in each condition, where ns denotes a nonsignificant result. **E** Neutrophil extracellular traps. Differentiated CD34^+^ cells were incubated with (right) or without (left) 50 nM PMA for 2 hours, stained with 0.2 *µ*M Sytox Green. Images are representative NETs from one of three biological replicates. Scale bars in these images show a distance of 10 *µ*m. **F**Neutrophil granule proteins. Differentiated CD34^+^ cells were deposited on glass slides, air dried, stained with antibodies to neutrophil elastase (NE, green) and myeloperoxidase (MPO, red), and nuclei counterstained with DAPI. Images are representative of stained cells from three distinct donors. Scale bars in these images show a distance of 2 *µ*m. All images were obtained on a Zeiss LSM 980 confocal microscope.

Another key function of neutrophils is the production of neutrophil extracellular traps (NETs), structures comprised of networks of DNA and granule components such as myeloperoxidase, which effect extracellular bacterial killing ^9^. The protein kinase C agonist phorbol 12-myristate 13-acetate (PMA) is a potent inducer of NET production (NETosis). Differentiated CD34^+^ cells incubated with 50 nM PMA for 2 hours produced NETs visualised as extracellular DNA seen when stained with Sytox Green (Fig. 2E). Intracellular staining for myeloperoxidase and neutrophil elastase showed strong staining for both granule proteins (Fig. 2F). Taken together, CD34^+^ HSPC-derived neutrophils were observed to recapitulate multiple functions of mature human neutrophils.

### 2.2 Lentiviral transduction and directed differentiation of HSPC produces transgene-expressing neutrophils

The expression of transgenes is an important tool for understanding the roles of proteins in health and disease. Mature neutrophils have an insufficient lifespan in culture to allow lentiviral transduction, which requires a minimum of 48 hours and results in cellular priming/activation. We therefore explored the feasibility of stable transduction of granulocytic precursors followed by directed differentiation. Several barriers to expressing transgenes in HSPC include difficulty of CD34^+^ HSPC transduction and temporal loss of transgene expression due to position-effect variegation resulting from random site vector integration. Fig. 3A shows the vector used in initial transduction experiments. We selected the fluorescent marker tagBFP due to its brightness and emission in the violet spectrum, freeing up the remainder of the visible spectrum for other fluorescent readouts. We found that a lentiviral transduction by spinfection with polybrene at a high multiplicity after 72 hours of cytokine stimulation yielded a high and uniformly transduced population of HSPC. Unexpectedly, we found that transgene expression is rapidly lost during G-CSF directed differentiation. To ameliorate this loss, we staggered transduction of differentiating progenitors in order to maintain expression. Cells were transduced at day 3, day 8, and day 13 of differentiation (Fig. 3B). During this period, cells were not exposed to lentivirus or polybrene for more than 48 hours, and media was changed at day 13 to prevent components of the transduction mixture from activating mature neutrophils. The final population contained around 40% transgene-expressing neutrophils with high expression of tagBFP, even on a modest PGK promoter (Fig. 3C,D). Thus, staggered transduction of granulocytic precursors during directed differentiation represents an effective approach to express transgenes in mature human neutrophils.

**Fig. 3.**
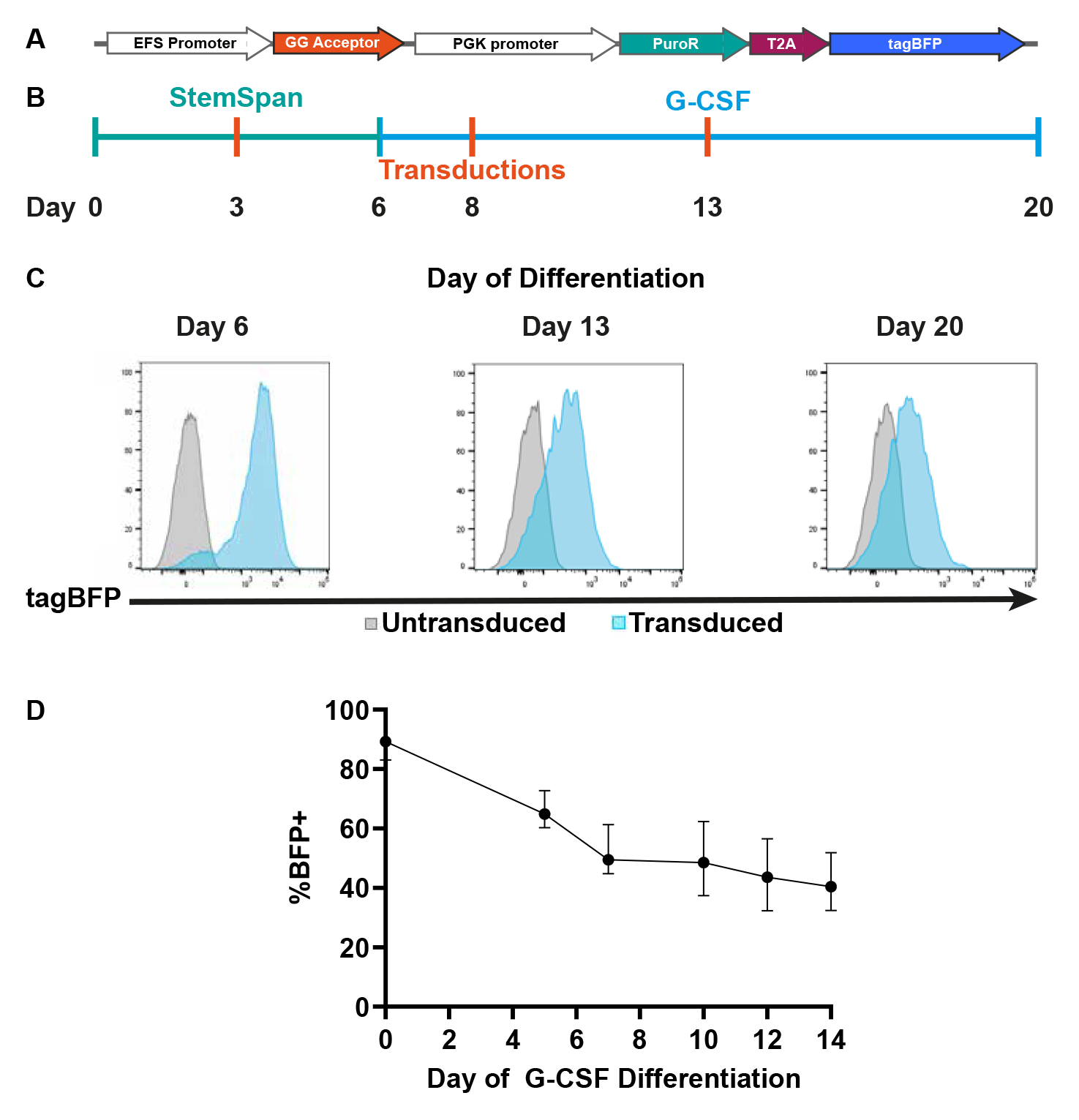
**A** Map of the lentiviral payload used in initial transduction optimisation experiments. The puro-T2A-tagBFP protein here is driven by a constitutive PGK promoter. Lentiviral elements up-and downstream of the payload are not shown. **B** Timeline of staggered lentiviral transduction of differentiating CD34^+^ HSPC. Progenitors were expanded in StemSpan for six days (teal), and then in G-CSF for 14 days (blue). Spinfection was carried out every five days from day three as shown in orange. **C** Representative histograms of tagBFP expression during differentiation at days six, 13, and 20. Experiments show live single cells whose gating is shown in Fig. S1C. **D** tagBFP expression in transduced cells during G-CSF directed differentiation. Live single cells were classed as tagBFP^+^ by drawing a gate on the untransduced population to a positivity of 0.5% (Fig. S1C). Data are shown as median (dot) and interquartile range (error bars) of twelve biological replicates. All data was obtained on an Attune NxT flow cytometer.

### 2.3 MicroRNA-driven RNA interference results in effective gene silencing in neutrophils

Silencing by RNA interference (RNAi) is an important tool to study functional contributions of genes of interest, but is presently impossible in primary human neutrophils due to their short lifespan and the toxicities inherent in the cationic delivery systems. However, our method of staggered lentiviral transduction of differentiating progenitors presents an avenue for delivery of RNA-i components for gene silencing prior to the acquisition of priming characteristics. To optimise the development of gene silencing systems we chose as our target *ITGAM*, the product of which is CD11b. *ITGAM* is highly expressed at mRNA and protein level in both myeloid progenitors and mature neutrophils. Surface CD11b is upregulated predictably between day 6-8 of G-CSF directed differentiation, allowing us to detect any disturbance of myeloid differentiation as a result of lentiviral or shRNA delivery. Additionally, cell surface CD11b is rapidly up-regulated in response to neutrophil activation, allowing simultaneous detection of off-target activation in the mature neutrophil population.

Alternate RNAi-like approaches using single stranded, RNAse-H activating antisense oligonucleotide (GapmeR) technologies have been shown to effect potent gene knockdown without the need for chemical delivery methods ^10^. However, high-dose, continuous GapmeR delivery failed to suppress *ITGAM* expression at mRNA and protein level in differentiating CD34 HSPC (Fig. S3). We next examined whether activating RNAi via stable expression of short hairpin RNA (shRNA) would result in efficient gene knockdown by constructing a bicistronic vector (Fig. 4A) expressing shRNA from a U6 promoter and selected in-assay by tagBFP expression. Nascent U6-driven shRNA constructs are transcribed as 21nt double-stranded RNA (dsRNA) hairpin species with a circular microRNA miR-21 derived loop, and terminating with a polyuridine tract that serves as the RNA polymerase III termination signal (Fig. 4B). When processed by DICER, these form short dsRNA species that can activate RNAi when loaded into the argonaute proteins. Using this system, we found that the the non-targeting sequence caused increased expression of CD11b in differentiated CD34^+^ cells. All three shRNA’s tested showed suppression of surface CD11b compared to the nontargeting shRNA, but not compared to the un-transduced control (Fig. 4C). This failure of silencing may be the result of CD11b upregulation by cellular priming/activation masking an effective RNAi-driven silencing response. shRNA hairpins transcribed by RNA polymerase III contain a 5’ triphosphate that is readily sensed by Retinoic Acid Inducible Gene-I (RIG-I) ^11^, a cytosolic RNA sensor which is known to activate neutrophils ^12^, and therefore approaches using shRNA driven by U6 and H1 promoters are likely to result in unacceptable off-target activation.

**Fig. 4.**
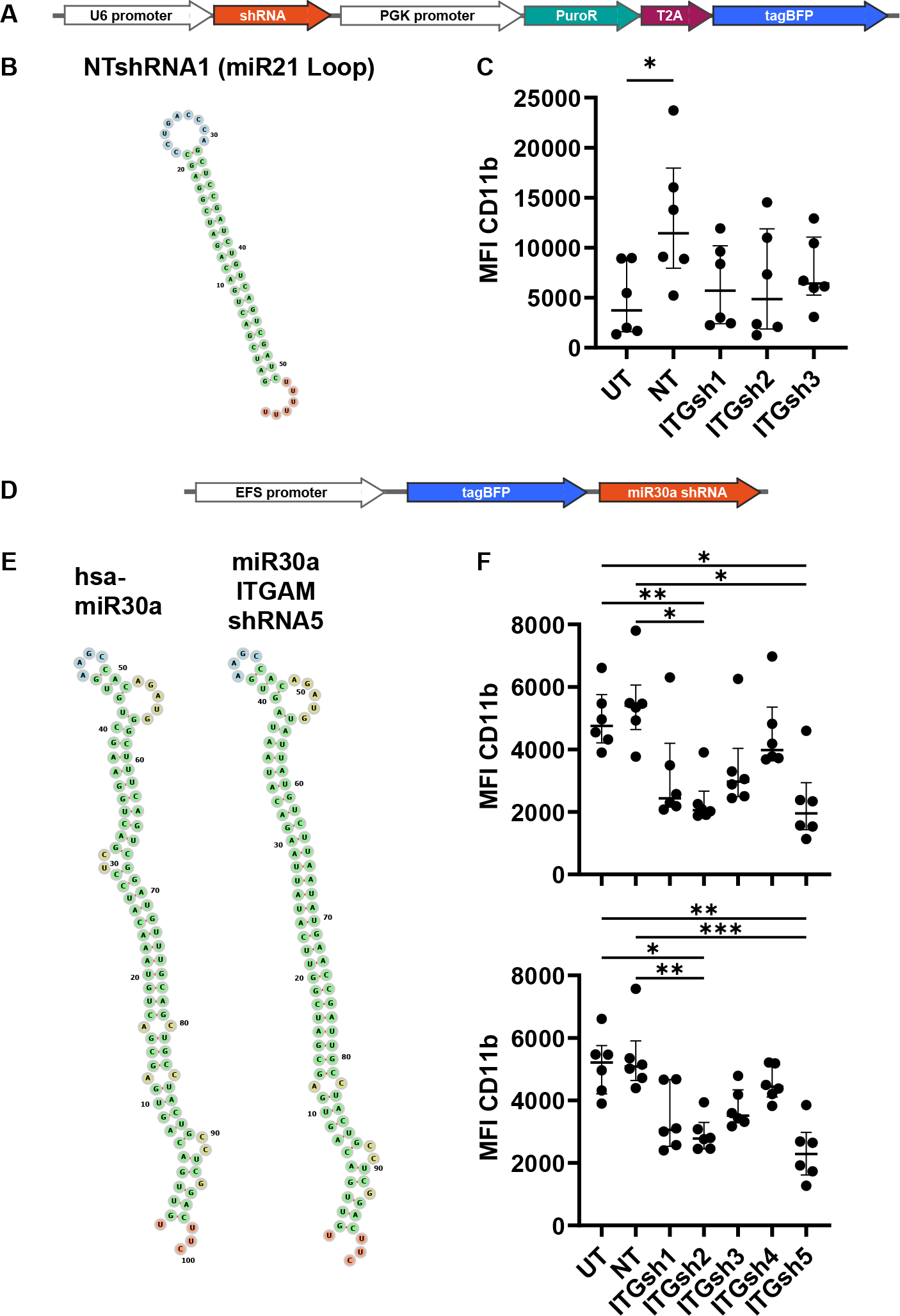
**A** Map of the payload for U6-shRNA vectors. This is a bicistronic construct containing U6-shRNA followed by PGK-Puro-T2A-BFP. **B** Secondary structure of U6-driven shRNA hairpin. The predicted transcript contains a 21nt dsRNA stretch separated by a 9nt hairpin derived from hsa-miR21. **C** CD11b expression assessed at day 20 of differentiation by flow cytometry. Cells underwent staggered transduction during differentiation with lentivirus carrying three different U6-driven shRNA’s targeting ITGAM, as well as a nontargetign control. **D** Map of the payload for the tagBFP-miRshRNA vector, with the shRNA in the 3’UTR of tagBFP. In all maps, lentiviral elements up-and downstream of the payload are not shown. **E** Secondary structures of endogenous hsa-miR-30a (left) and miR30a-ITGAM-sh5 which contains a guide sequence targeted to the 3’LTR of *ITGAM*. The structure of the synthetic shRNA differs from endogenous in missing two bulges: a 1nt mismatch at position 12, and a 2nt mismatch at positions 30-32. All structures were generated with the RNAfold Webtool ^16^. **F** Experiments showing CD11b expression by flow cytometry at day 20 of differentiation after staggered transduction with lentivirus carrying miR-shRNA’s. Five different miR-shRNA’s targeted to *ITGAM* as well as a nontargeting control were tested. Data for both tagBFP^+^ cells (top) or all live single cells (bottom) are shown. UT, untreated; NT, nontargeting shRNA; ITGsh1-5, shRNA’s 1-5 targeted to *ITGAM*. For figures 4C and 4F, individual data from six distinct donors are shown as dots, with horizontal bars indicating the median of each group with the interquartile range depicted by error bars. Asterisks indicate statistical significance (Friedman’s test) as detailed by bars between relevant groups: *p*≤*0.05;**p*≤*0.01; ***p*≤*0.001. Gating strategy for all flow cytometry is shown in Fig. S1C, and was obtained on the Attune NxT flow cytometer.

Endogenous effectors of RNAi are known as micro RNA’s (miR’s), noncoding sequences that form complex hairpins that are sequentially processed by the conserved microprocessor (DROSHA/DGCR8) and DICER enzymes to yield 22-nucleotide RNA duplexes with 2 nucleotide 3’ overhangs, which load into the RNA-induced silencing (RISC) effector complex ^13^. Modification of the 3p arm of well characterised microRNA’s such as miR-30a can result in potent suppression of target genes ^14, 15^. Additionally, the dsRNA species generated from miR-shRNA’s lack the activating 5’ triphosphate seen in RNA polylmerase III transcribed shRNA’s, and we predicted these would be capable of activation-free gene silencing. We designed a lentiviral vector containing an EF1*α*-promoter driven tagBFP construct with a miR-30a scaffold in the 3’ untranslated region capable of accepting target shRNA sequences (Fig. 4D). The predicted secondary structures of the microRNA-platformed shRNA’s mimicked that of the endogenous hsa-miR30a, differing only by two bulges present in the native stem-loop sequence (Fig 4E). Two of the five shRNA’s tested significantly suppressed surface CD11b levels without significant up-regulation in the non-targeting control. Importantly, this effect was seen both in tagBFP^+^ (Fig. 4F, top) and the entire population of differentiated CD34^+^ cells at day 14 (Fig. 4F, bottom). This may reflect the high level of transduction present during directed differentiation, as well as a prolonged RNAi-induced silencing effect even in differentiated cells that have lost tagBFP-miRshRNA expression. Taken together, miR-shRNA induction of RNAi in differentiating progenitors is capable of potent, activation-free gene silencing in differentiated neutrophils.

### 2.4 Cas9-gRNP genome editing results in high-efficiency gene knockout in neutrophils

We next examined the effectiveness of gene knockout in neutrophils using CRISPR-Cas9 technology. Efficient delivery of Cas9 complexed with guide RNA (Cas9-gRNP) has been described using nucleofection of CD34^+^ HSPC ^8^, and we used this approach to knockout *ITGAM*. Cas9-gRNP complexes were nucleofected into HSPC after prestimulation with early-acting cytokines, and cells were then directed towards neutrophil differentiation after a 72-hour resting period. The timeline of nucleofection is shown in Fig. 5A. We initially tested three separate guide sequences (Fig. S4) targeting early exons in the *ITGAM* gene in order to increase the chance of significant mutations prior to the membrane-spanning domain.

**Fig. 5.**
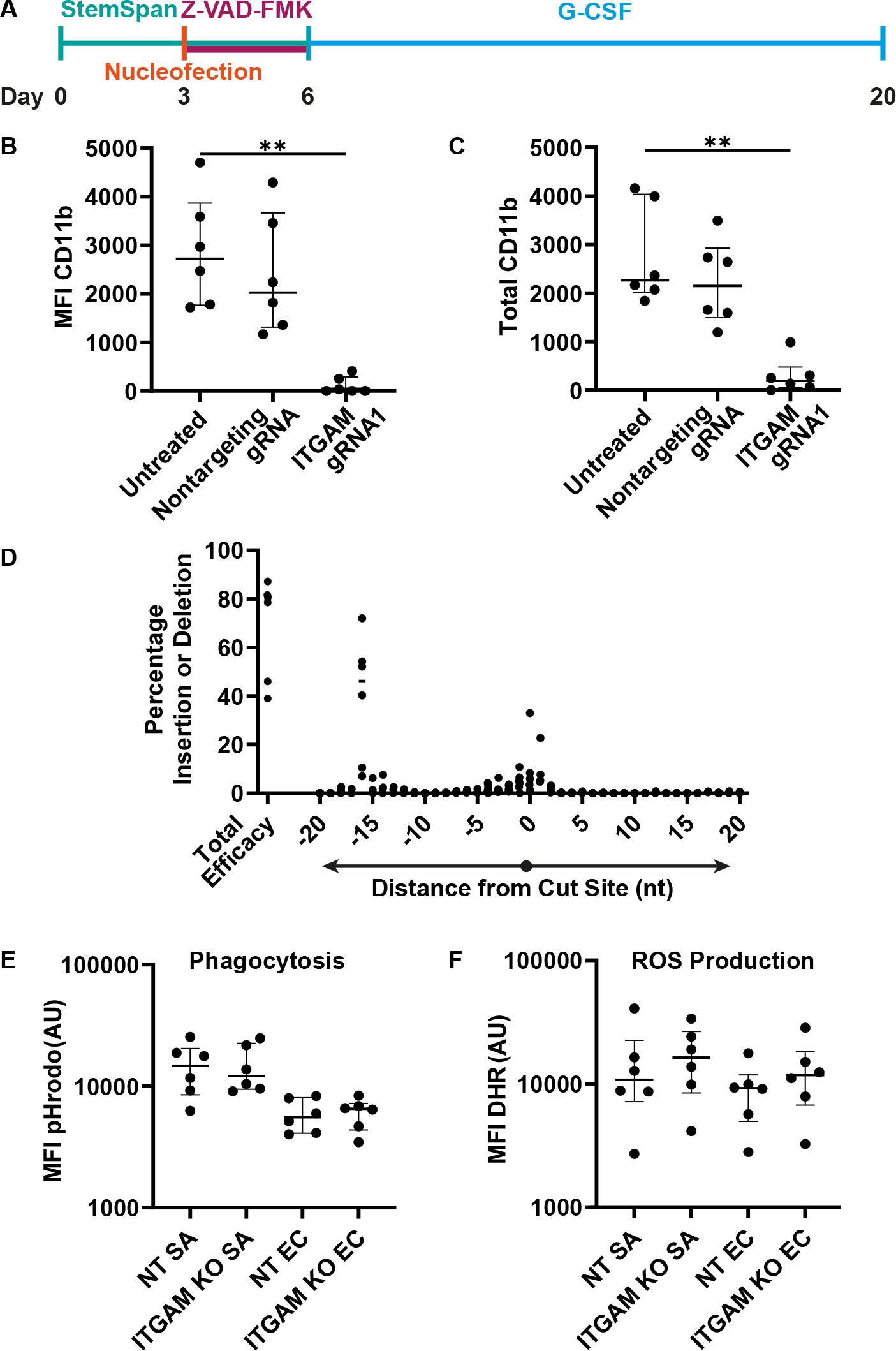
**A** Differentiation timeline with nucleofection (red) after 3 days of StemSpan preconditioning. Nucleofected cells were rested in media containing StemSpan and 5 *µ*M Z-VAD-FMK, a pan-caspase inhibitor, to mitigate cell losses after nucleofection. Cells were then differentiated G-CSF at 10ng/mL for 14 days. **B,C** CD11b expression by flow cytometry at day 20. MFI (B), and total CD11b - the product of percent positive and MFI of the positive cells (C) are shown. **D**Genomic indel profiles obtained by Sanger sequencing and Tracking of Indels by DEcomposition (TIDE) analysis comparing *ITGAM* guide RNA 1 compared to donor-specific nontargeting controls. The Cas9 cut site is shown as 0nt, and insertions (right, positive) and deletions (left, negative) are expressed as a percentage of total sequences. Data at 0nt are indicate no sequence change. Total efficacy is shown at the left of the graph and is the sum of percentage indels. **E,F** Functional capacities of CRISPR-edited neutrophils. Phagocytosis of serum opsonised pHrodo red conjugated bioparticles of *S. aureus* (SA) and *E. coli* (EC) are shown in (E), and oxidative response by DHR fluorescence are shown in (F). NT, Nontargeting guide; ITGAM KO, ITGAM gRNA1. For figures 5B-C and 5E-F, individual data from six distinct donors are shown as dots, with horizontal bars indicating the median of each group with the interquartile range depicted by error bars. Asterisks indicate statistical significance (Friedman’s test) as detailed by bars between relevant groups: *p*≤*0.05;**p*≤*0.01; ***p*≤*0.001. Gating strategies are shown in Fig. S1D and all flow data was obtained on the Attune NxT flow cytometer. Example outputs of TIDE analysis are shown in Fig.S2.

ITGAMgRNA1-Cas9 complexes electroporated into CD34^+^ HSPC were able to effect potent knockdown of *ITGAM* in the entire differentiated CD34^+^ cell population at the protein (Fig. 5B-C) and genomic level (Fig. 5D). CRISPR knockout by Cas9-gRNP nucleofection relies heavily on nonhomologous end joining to repair double strand breaks, which typically results in random indel profiles around the breakpoint. In four of the six donors, however, a 16nt deletion was the predominant mutation accompanying the break-point proximal indel profile, which may be the result of microhomology-mediated end joining seen in a small proportion of gRNA species ^17^. This mutation is predicted to result in an early premature stop codon in Exon 3 of *ITGAM* which is proximal to the membrane spanning domain (Fig. S2C).

We chose this method to investigate the contribution of CD11b to important neutrophil functions. The product of the *ITGAM* gene, CD11b, is also known as integrin *α*M, an integrin which complexes with CD18 to form the C3bi complement receptor CR3, where CD11b contains the important *α_M_ I* domain ^18^. CR3-C3bi interactions are thought to be important in the phagocytosis of complement-opsonised pathogens. Surprisingly, CRISPR-Cas9 mediated knockdown of *ITGAM* did not have a significant impact on phagocytosis of serum-opsonised *S. aureus* or *E. coli* bioparticles *in vitro* (Fig. 5E), and did not affect the resultant oxidative response (Fig. 5F), suggesting that in whole serum opsonised bacteria, the contribution of CR3-C3bi may be dispensable. Here we have shown that our model of Cas9-RNP genome editing and directed differentiation is capable of efficient knockdown of highly expressed genes to examine their contribution to important phenotypic functions in human neutrophils.

## 3 Methods

### 3.1 Magnetic isolation of CD34^+^ HSPC

Apheresis cones are used to deplete leukocytes from blood during platelet donation, and are a byproduct of this process. Each cone contains approximately 1 *×* 10^9^ peripheral blood mononuclear cells. Cones were obtained from NHS Blood and Transplant and were a kind gift from Dr Mattia Frontini. Blood was eluted from cones by washing with Dulbecco’s phosphate buffered saline (dPBS) containing 0.38% trisodium citrate (HiMedia), layered onto Lymphoprep (StemCell Technologies) density gradient medium, centrifuged at room temperature for 30 minutes at 800g, and peripheral blood mononuclear cells were removed from the interface. CD34^+^ HSPC were positively selected using the EasySep^TM^ Human CD34 Positive Selection Kit II (StemCell Technologies) according to manufacturer’s instructions. Isolated CD34^+^ HSPC were cryopreserved in freezing medium containing 50% Foetal Bovine Serum (Gibco), 40% SFEM II (StemCell Technologies) and 10% DMSO (Sigma-Aldrich).

### 3.2 CD34^+^ HSPC expansion and granulocytic differentiation

CD34^+^ HSPC were defrosted with routinely over 90% viability. Cells were plated at a minimum density of 5 *×* 10^5^ cells/mL in SFEM II medium (StemCell Technologies) with StemSpan^TM^ CD34+ Expansion Supplement (StemCell Technologies). Cells were incubated at 37°C with 5% CO_2_. On day 6, medium was changed to SFEM II containing 10ng/mL recombinant human granulocyte colony stimulating factor (StemCell Technologies) and maintained in this medium until day 20, with medium being changed every 48 hours, and where cell density did not exceed 1 *×* 10^6^ cells/mL. At day 20, cells were washed with dPBS and phenotyped.

**3.3 Isolation of human peripheral blood neutrophils**

Human peripheral blood neutrophils were obtained from consenting healthy volunteers using discontinuous Percoll (GE Healthcare) density gradient centrifugation. 40 mL of whole blood was anticoagulated with trisodium citrate at a final concentration 0.38% and centrifuged at 300g for 20 minutes to pellet all blood cells. 10 mL of plasma was converted into autologous serum by addition of 0.22mmol CaCl_2_ and incubation at 37°C in glass vials for 30 minutes. The remainder of the plasma was clarified by centrifugation at 1400g for 20 minutes to remove platelets. Red cells were sedimented in 0.75% dextran T-500 (Pharmacosmos) for 30 minutes, and the remaining upper mixed leukocyte layer was pelleted, resuspended in autologous clarified plasma, and underlaid with a discontinuous gradient comprising 2mL each of 42% and then 51% Percoll. This was centrifuged for 14 minutes at 150g with a slow acceleration and no brake. Neutrophils were removed from the 42%/51% interface, washed three times in dPBS by centrifugation at 300g, and used as described in downstream phenotyping assays.

### 3.4 Cytospin preparation and staining

Cells (2 *−* 5 *×* 10^5^) were placed in a cytofunnel and centrifuged at 300g onto polylysine-coated glass slides using a Cytospin 3 cytocentrifuge (Shandon). Slides were air dried, fixed for 4 minutes in methanol, and stained using the DIFF-QUIK (RAL Diagnostics) modified Romanowsky stain. Slides were immersed in a buffered Eosin Y solution (DIFF-QUIK red) for 1 minute and then counterstained in a solution of methylene blue and Azure A (DIFF-QUIK blue) for 3 minutes. These were washed gently with water and mounted with DPX histomountant (Sigma-Aldrich). Microscopic images were obtained using an ECHO Revolve microscope.

### 3.5 Staining and flow cytometric analysis

Cells were stained at a final concentration of 5*×*10^5^ cells/mL in staining buffer (dPBS with 1% bovine serum albumin) for 30 minutes at room temperature) with fluorescent conjugated antibodies. Stained cells were analysed on the following machines as indicated in the text: LSR Fortessa(Beckton Dickinson) or Attune NxT (Thermofisher Scientific). Representative gating strategies are shown in the relevant supplementary Fig. S1A-D. All analysis was carried out on between 10,000-20,000 live single cells unless otherwise specified. The following antibodies or staining reagents were used: Mouse-anti human CD11b-FITC (Bear-1, Hycult Biotech), mouse-anti human CD62L-Alexa Fluor 647 (DREG-56, Biolegend), mouse-anti human CD16-Pacific blue (3G8, Biolegend), mouse anti-human CD11b-Alexa Fluor 647 (ICRF44, Biolegend), Annexin V-FITC (Thermofisher Scientific), propidium iodide (Biolegend).

### 3.6 Priming and chemiluminescent reactive oxygen species (ROS) detection

Extracellular ROS was assayed via the luminol-HRP system. Luminol is oxidised in a chemiluminescent reaction to 3-aminophthalate by HRP in the presence of peroxide radicals. Neutrophils or differentiated CD34^+^ cells were primed with 1 *µ*M platelet activating factor (Sigma-Aldrich) or dPBS vehicle, and incubated in a Thermomixer Comfort shaking heat block (Eppendorf) at 37°C for 5 minutes. Luminol (Sigma-Aldrich) was added to a final concentration of 1 *µ*M, incubated for 3 minutes, and cells were pipetted into individual wells of a 96-well white opaque optical plate pre-loaded with 50 units of HRP (Sigma-Aldrich). This was introduced into a warmed (37°C) luminometer (Berthold LB 960) which has an inbuilt injector function. For experiments with fMLP stimulation, fMLP (Sigma-Aldrich) was injected to a final concentration of 100 nM. Luminescence was measured thereafter at 5 second intervals for 10 minutes to give a dynamic time course.

### 3.7 Flow cytometric quantitation of phagocytosis

For all experiments, neutrophils or differentiated CD34^+^ cells were suspended in RPMI 1640 (Thermofisher) with 10 mM HEPES (Gibco) at 37°C. pHrodo-red labelled *Staphylococcus aureus* or *Escherichia coli* bioparticles (Invitrogen) were serum opsonised for 30 minutes in 50% human serum and then used in-assay at a concentration of 25 *µ*g/mL. 5*×*10^5^ cells were incubated in 2 mL round bottom Eppendorf tubes at a final volume of 250 *µ*L with serum-opsonised pHrodo red-conjugated bioparticles and dihydrorhodamine (Invitrogen, 3 *µ*M) for 30 minutes at 37°C on a Thermomixer Comfort shaking heat block (Eppendorf). Simultaneous identical cold control experiments were performed on ice for 30 minutes. Phagocytosis was halted by quenching tubes on ice for 10 minutes while staining with propidium iodide (Biolegend) and mouse anti-human CD16-Pacific blue antibody (3G8, Biolegend), and analysed on the LSR Fortessa II (Beckton Dickinson) or Attune NxT (Invitrogen) flow cytometers. Live CD16^+^ cells were selected for analysis and assessed for pHrodo-red and dihydrorhodamine fluorescence.

### 3.8 Confocal imaging of intracellular proteins and neutrophil extracellular traps

Differentiated CD34^+^ cells were deposited on polylysine slides as described above. Slides were air dried and then fixed with 4% paraformaldehyde for 15 minutes at room temperature. Cells were block-permeabilised with 5% BSA and 0.1% Triton X-100 in dPBS for 30 minutes at room temperature. Slides were washed in dPBS three times and stained with primary antibodies: goat anti-human neutrophil elastase (Santa Cruz Biotech 9520, 1:1000), and goat anti-human myeloperoxidase (Abcam 9535) in 5% BSA at 4°C overnight.

They were then washed with dPBS and stained for 1 hour at room temperature with donkey anti-goat Alexa-Fluor 488 secondary antibody (Invitrogen A11055, 1:400), washed again, and then subsequently stained with goat anti-rabbit Alexa-Fluor 568 (Invitrogen A11011) in 5% BSA for 1 hour at room temperature. Slides were washed after secondary staining and mounted with ProLong Gold Antifade Mountant with DAPI (Life Technologies). To assess NETosis, differentiated CD34^+^ cells were resuspended in Dulbecco’s dPBS with or without 50 nM PMA, and deposited into a 1cm^2^ area delineated by water repellent ink. Cells were placed in a humidified chamber and incubated for 2 hours at 37°C and 5% CO_2_. Media was gently aspirated and cells stained with dPBS containing 0.2 *µ*M of Sytox Green (Invitrogen) at room temperature for 15 minutes. These were washed gently and fixed in methanol for 4 minutes, air dried, and mounted with DPX mountant for histology. All imaging was performed on a Zeiss LSM 980 confocal microscope (Carl Zeiss AG).

### 3.9 RNA isolation and qPCR

Cells were lysed in RLT buffer (Qiagen) supplemented with 1% ß-mercaptoethanol (*β*-ME) and homogenised with a syringe and needle by aspirating 10 times. RNA was extracted using RNeasy Micro Kit (Qiagen) with the on-column DNase digestion step, eluted in 30 *µ*L of RNAse-free water and measured the concentration on a Nanodrop Lite Spectrophotometer (ThermoFisher Scientific). cDNA was prepared using the High Capacity cDNA Reverse Transcription Kit (Applied Biosystems) according to the manufacturer’s instructions. qPCR reactions were prepared using SYBR green PCR Jumpstart Taq ReadyMix (Sigma-Aldrich), ROX Reference Dye (ThermoFisher Scientific, US), 0.4 *µ*M primers (Qiagen), and cDNA to a total input volume of 10 *µ*L. Reactions were analysed in triplicate on a QuantStudio 6 Flex (Applied Biosystems) PCR machine. Gene expression was normalised to the geometric mean of three reference genes (B2M, GNB24 and RPL32), and the fold differential expression was determined with the 2*^−^*^ΔΔ*Ct*^ method.

### 3.10 Plasmids and Transfer Vector Construction

All plasmids used in this work are credited in supplementary Fig. S8. Detailed plasmid maps and construction are shown in fig.S5. Transfer vectors were produced using Gibson Assembly ^19^, a cloning technique which seamlessly joins dsDNA fragments with overlapping sequences at their termini. Primers can be designed to incorporate these overlaps to allow a ‘copy-and-paste’ construction of desired transfer plasmids. Detailed construction of these vectors, Gibson Assembly primers, and maps are shown in supplementary Fig. S6. Individual segments were amplified using primers at a concentration of 0.5 *µ*M with Platinum SuperfiII polymerase (Invitrogen) and 35 cycles according to manufacturers’ instructions for extension time, with a final extension of 10 minutes. Size was validated by agarose gel electrophoresis and PCR products were purified using QiaQuick PCR Purification Kit (Qiagen). Equimolar amounts of purified DNA fragments (minimum 15fmol) were incubated at 60°C for 1 hour with a Gibson Mastermix to achieve a final concentration of the following: 4U/mL T5 exonuclease, 25U/mL Phusion polymerase, 4000U/mL Taq ligase, 10mM Mg++, 5% PEG-8000, 10mM Tris-HCl, 10mM DTT, 200µM dNTP. All enzymes were obtained from New England Biolabs. 1 *µ*L of crude reaction was used to transform NEB Stable competent cells by heat shock according to manufacturers’ instructions and grown at 32°C overnight on Luria-Bertani agar containing ampicillin (Gibco) at 100 *mu*g/mL. Plasmids from individual colonies were sequenced by Sanger Sequencing (Azenta, UK).

### 3.11 Lentiviral vector production

Lentiviral particles were produced in HEK293T packaging cells seeded at a density of 60,000 cells/cm^2^ 24 hours prior to transfection, and pretreated with 25 *µ*M chloroquine in fresh complete medium for six hours on the day of transfection. A packaging mixture of 23 fmol/cm2 of PSPAX2 and 3.3 fmol/cm2 of pMD2.G were used with 30 fmol/cm2 of the appropriate transfer plasmid. Plasmids were mixed in warmed Opti-MEM (Gibco) with linear polyethyleneimine (PEI 25kDa, Merck) at a PEI:DNA ratio of 3:1, incubated for 15 minutes at room temperature, and added to cells. Supernatants were harvested every 24 hours for 3 days, pooled, clarified by centrifugation at 4000g for 15 minutes, and lentiviruses pelleted by ultracentrifugation at 25000rpm for 2 hours at 4°C in a Beckman-Coulter XPN-100 centrifuge using a Sw-32 Ti rotor. Pellets were resuspended in SFEM II, snap frozen, and titred on HeLa cells by logarithmic dilution in the presence of 4 *µ*g/mL polybrene. Where large volumes of supernatant were used, precipitation in 10% PEG-8000 and 0.15M NaCl at 4°C overnight prior to ultracentrifugation.

### 3.12 Cas9-gRNP complexing and nucleofection

Alt-R SpCas9 V3 nuclease, crRNA, tracrRNA, and electroporation enhancer were all obtained from Integrated DNA Technologies. crRNA sequences for each gene were designed using the CRISPOR tool ^20^ and picked according to the highest Moreno-Mateos score and lowest off-target scores. gRNA complexes were formed by annealing equimolar (1 pmol) amounts of crRNA and tracrRNA by heating to 95°C for 5 minutes and cooling on benchtop to room temperature. These were complexed with 0.4pmol Cas9 for 15 minutes at room temperature and then used immediately in electroporation reactions. CD34^+^ HSPC selected as previously described were preconditioned in SFEM II and StemSpan cytokines for 72 hours and electroporated with the Cas9-gRNP mixture using the Human CD34^+^ Cell Nucleofector kit (Lonza) in a Lonza IIb Nucleofector using program U-008, according to manufacturers’ instructions, in the presence of 4 *µ*M of Electroporation Enhancer. Cells were rested in SFEM II containing StemSpan cytokines and 5 *µ*M Z-VAD-FMK (ApexBio) for 72 hours and then directed to granulocytic differentiation as described above. Knockout efficacy was validated by flow cytometry and by targeted sequencing. Mutational profile at the gRNA target site was assessed using TIDE analysis ^21^ shown in Fig. S2. Briefly, genomic DNA was isolated from day 14 differentiated CD34^+^ cells using the Purelink Genomic DNA kit according to manufacturer’s instructions.

Genomic regions flanking the target site for ITGAMgRNA1 were amplified using primers AGGAAGACCTTTTCTCATTCTGAGT (forward) and GCTTCCTGTCCTTGTACCCC (reverse). Amplification conditions were as follows: Primers at 0.5 *µ*M, Superfi II Platinum Enzyme (Invitrogen), input genomic DNA 100ng, volume 50 *µ*L, 45 cycles, extension time 1 minute, final extension 5 minutes. Size checked by gel electrophoresis and products were purified with the QiaQuick PCR Purification Kit (Qiagen) prior to Sanger Sequencing. Raw sequence files are entered into the TIDE decomposition tool ^21^, found at http://shinyapps.datacurators.nl/tide/. Guide sequences are shown in Appendix A.8.

### Ethical approval declarations

Apheresis cones used in this study were obtained after written consent from individual blood donors under NHS Blood and Transplant Guidelines. Informed, written consent was obtained from donors for experiments involving peripheral blood samples, and the methods were given favourable ethical approval by the Cambridge Central Research Ethics Committee (ref 06/Q0108/281).

## 4 Discussion

Here, we have presented methods for efficient gene manipulation of neutrophils differentiated from CD34^+^ HSPC. The differentiated cells recapitulate key neutrophil functions such as the ability to prime their respiratory burst to formyl peptides, and phagocytosis. This method is rapid, scalable, and more cost-effective than animal models. Importantly, magnetic isolation of CD34^+^ HSPC is feasible from patient samples, and which will allow patient-and disease-specific study of neutrophil biology. Dissecting the roles of specific genes is critical to understanding diseases in which neutrophils become dysfunctional. The mechanisms that result in such dysfunction remain poorly characterised, and the ability to genetically manipulate neutrophils will unlock new insights into the biology of neutrophils in health and disease.

There are a number of limitations of both the *ex vivo* differentiation protocol and its genetic manipulation. Although a high proportion of neutrophils in the day 14 population are segmented, the presence of progenitors and other myeloid cells may interfere with functional assays. Although this can be mitigated by in-assay selection of mature neutrophils using CD16, in chromogenic or chemiluminescent bulk assays immature neutrophils and their precursors may dilute important effects. This can be seen in the lower magnitude of the respiratory burst to formyl peptides in the priming assay. It is possible to magnetically enrich mature neutrophils by negative selection ^22^, but this may result in cell loss or unintended activation if not performed with the utmost care. Additionally, it is likely that even segmented cells are not fully mature, as their expression of key markers such as CD16 is much lower than isolated circulating neutrophils. This is not for a lack of G-CSF receptor signalling, as the concentrations of G-CSF used in culture exceed normal endogenous circulating concentrations of 100-1000 pg/mL ^23^, but rather may be due to the lack of mechanical and cellular components of the bone marrow microenvironment, or other complementary cytokines such as IL-6, IL-3, and GM-CSF. The exclusion of other cytokines in this method aids the generation of a highly pure population of neutrophils, as the inclusion of other cytokines may lead to non-neutrophil lineage commitment ^24^.

Although lentiviral transgene delivery is efficient and immunologically silent ^25^, we have unexpectedly found loss of transgene expression during directed differentiation, the cause of which is unclear. This may be due to inevitable large-scale changes in chromatin availability which are a central feature of granulopoiesis ^26^. Lentiviral integration favours transcriptionally active chromatin such as that marked by histone H3 trimethylated at lysine 36 (H3K36me3) due to the importance of lens epithelial derived growth factor LEDGF(p75)-H3K36me3 interactions in lentiviral integration ^27^. H3K36me3 distribution changes dramatically during granulopoiesis ^26^, and it is likely that proviral transgenes delivered to early progenitors will be silenced by the large-scale chromatin changes that occur in differentiating cells. We have also observed stabilisation of lentiviral transgene expression after myeloid commitment as defined by the acquisition of CD11b at day 13 of differentiation (day 7 of G-CSF directed differentiation, Fig. 3D) which would support this mechanism. Despite these difficulties, staggered transduction still results in robust transgene expression in the final population of neutrophils. We have also shown that Cas9-gRNP mediated gene editing is possible to a high efficiency in primary human haematopoietic progenitors that results in activation-neutral gene knockdown in differentiated neutrophils that is reproducible across multiple donors. The efficient gene disruption seen in the bulk population of neutrophils obviates the need for cell sorting with attendant cell loss and activation. Our data shows that Cas9-gRNP delivery can also result in a high degree of precise edits.

The combined application of these novel approaches presents exciting new possibilities to the study of neutrophil biology which are core features of CRISPR-Cas9 technology: Base editing, saturating mutagenesis, and editing by homology-directed repair. Finally, the technological intersection of activation-neutral genome editing, transgene delivery, and reliable *ex vivo* granulopoiesis make previously unachievable forward genetic screens possible in human neutrophils, which have the potential to discover unknown facets of their biology, and presents exciting avenues for further research.

## Supplementary information

## Acknowledgments

Flow cytometry was supported by the Cambridge NIHR BRC Cell Phenotyping Hub who maintain the central phenotyping core facility but did not contribute intellectually to the design of this work.

## Declarations

- Funding AYKCN was supported by a Wellcome Trust PhD Fellowship Grant (222919/Z/21/Z). CS is supported by UKRI/Medical Research Council (MR/S035753/1, MR/X005070/1, MR/P502091/1), the National Institute for Health and Care Research (NIHR) (NIHR133788) and the NIHR Cambridge Biomedical Research Centre (BRC-1215-20014).
- Competing interests The authors declare no competing interests.
- Ethics approval Apheresis cones used in this work were obtained from anonymous, healthy blood donors who gave written consent for their use in research where the identity of the donor would not be traceable, under NHS blood and transplant guidelines. Research involving human peripheral blood received favourable ethical approval from the Cambridge Central Research Ethics Committee (ref 06/Q0108/281).
- Consent to participate - all healthy volunteers providing blood samples provided written informed consent in accordance with the ethical approvals in place.
- Consent for publication - Not applicable
- Availability of data and materials All plasmids generated in this work have been deposited with Addgene.
- Code availability - Not applicable.
- Authors’ contributions - A.Y.K.C.N and C.S. conceived the project, analyzed the data, wrote the manuscript; A.Y.K.C.N. developed the model and performed lentiviral and Cas9-gRNP experiments, J.B. helped with Cas9-gRNP optimisation and guide selection, M.F. provided the apheresis cones, and E.F. performed the GapmeR-knockdown experiments.

