## Supplemental data and figures. for "Efficient Methods for Target Gene Manipulation in Haematopoietic Stem Cell Derived Human Neutrophils"

### Appendix A Extended Data

#### A.1 Representative Gating Strategies

**Figure S1A. Gating for live/dead and marker analysis** Stained cells were analysed on the LSR Fortessa analyser (Beckton Dickinson). A polygon gate was drawn over the main cell population on a forward scatter area (FSC-A)/side scatter area (SSC-A) plot. Singlets were selected by gating a uniform area-width ratio. A quadrant gate on Annexin V-FITC (blue Laser 530/30) and propidium iodide (blue laser 695/40) all singlets was used to define live, apoptotic, and necrotic cells. For the analysis of cell surface markers CD11b, CD62L, and CD16, live cells were selected by propidium iodide exclusion. Marker positivity was defined by gating against isotype to a positivity of 0.5%.

**Figure S1B. Gating strategy for phagocytosis experiments** Stained cells were analysed on the LSR Fortessa analyser (Beckton Dickinson). A polygon gate was drawn over the main cell population on a forward scatter area (FSC-A)/side scatter area (SSC-A) plot. Live singlets were selected by gating a uniform area-width ratio and propidium iodide exclusion. ROS production was read in the blue laser 530/30 channel, and phagocytosis in the yellow laser 582/15 channel in all live singlets.

**Figure S1C. Gating for marker analysis on BFP$^+$ cells** Stained cells were analysed on the Attune NxT flow cytometer (Thermo Fisher Scientific). A polygon gate was drawn over the main cell population on a forward scatter area (FSC-A)/side scatter area (SSC-A) plot. Live singlets were selected by gating a uniform area-width ratio and propidium iodide exclusion. BFP$^+$ positive cells were defined by gating against untransduced cells to a positivity of 0.5% in the violet laser 450/50 channel.

**Figure S1D. Gating strategy for CRISPR-Cas9 experiments**

Stained cells were analysed on the Attune NxT flow cytometer (Thermo Fisher Scientific). A polygon gate was drawn over the main cell population on a forward scatter area (FSC-A)/side scatter area (SSC-A) plot. Live singlets were selected by gating a uniform area-width ratio and propidium iodide exclusion. $CD11b^+$ positive cells were defined by gating against untransduced cells to a positivity of 0.5% in the blue laser 530/30 channel.

#### Supplementary Figure 1

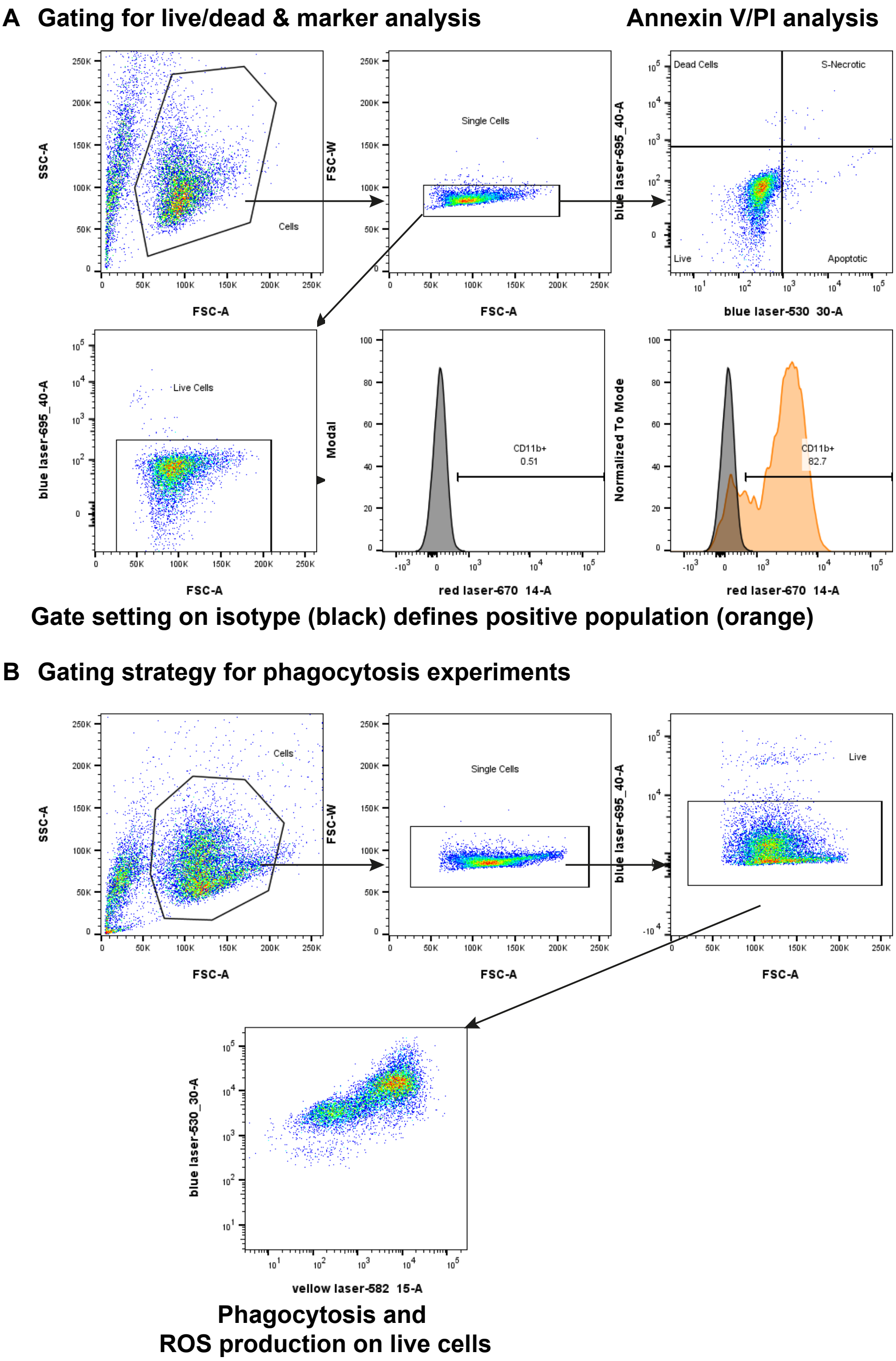

**Supplementary Figure 1**

#### C Gating for marker analysis on BFP$^{+}$ cells

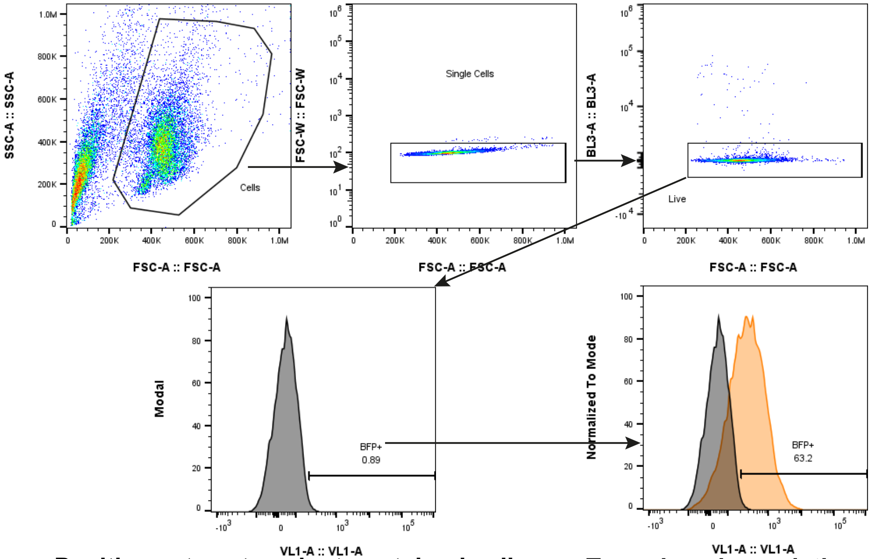


**Positive gate set against unstained cells (black)**

**Transduced population (orange)**

#### D Gating Strategy for CRISPR-Cas9 Experiments

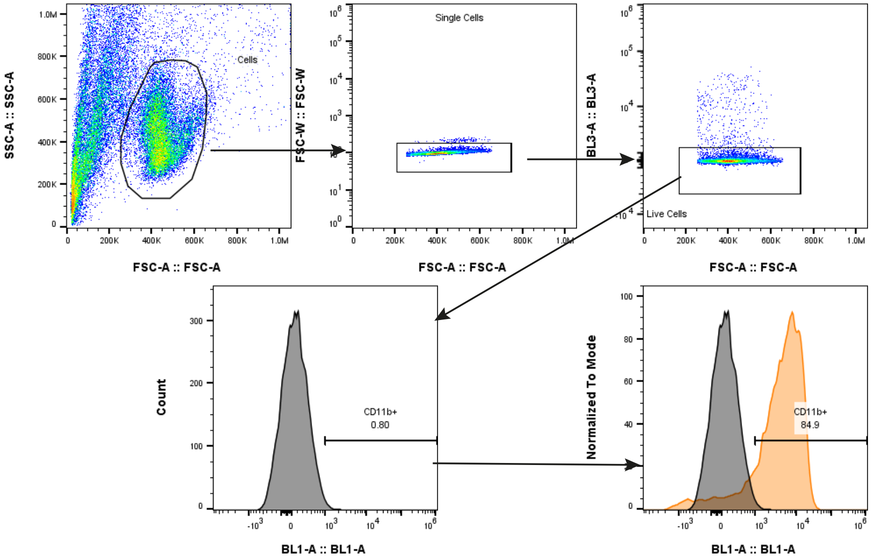


**Gate Setting on isotype (black) defines CD11b$^{+}$ population (orange)**

L

#### A.2 TIDE Analysis

Tracking of Indels by DEcomposition (TIDE) analysis was performed by uploading .ab1 trace files obtained from sanger sequencing onto the TIDE webtool at https://tide.nki.nl/. Genomic DNA amplified and sequenced from nontargeting-gRNA electroporated cells were used as the control sequences, and sample DNA was from Cas9-ITGAM-gRNA1 electroporated cells. When the guide sequence is provided to the webtool, it displays a plot of aberrant sequence fraction (green) around and after the expected cut site (Fig. S2A). This can be expressed as a proportion of insertions and deletions (Fig. S2B. A prevalent 16bp deletion seen in four of the six donors is shown here, alongside the breakpoint-proximal insertion-deletion pattern characteristic of nonhomologous end joining. The product of this 16bp deletion is a premature truncation product seen in Fig. S2C.

#### Supplementary Figure 2

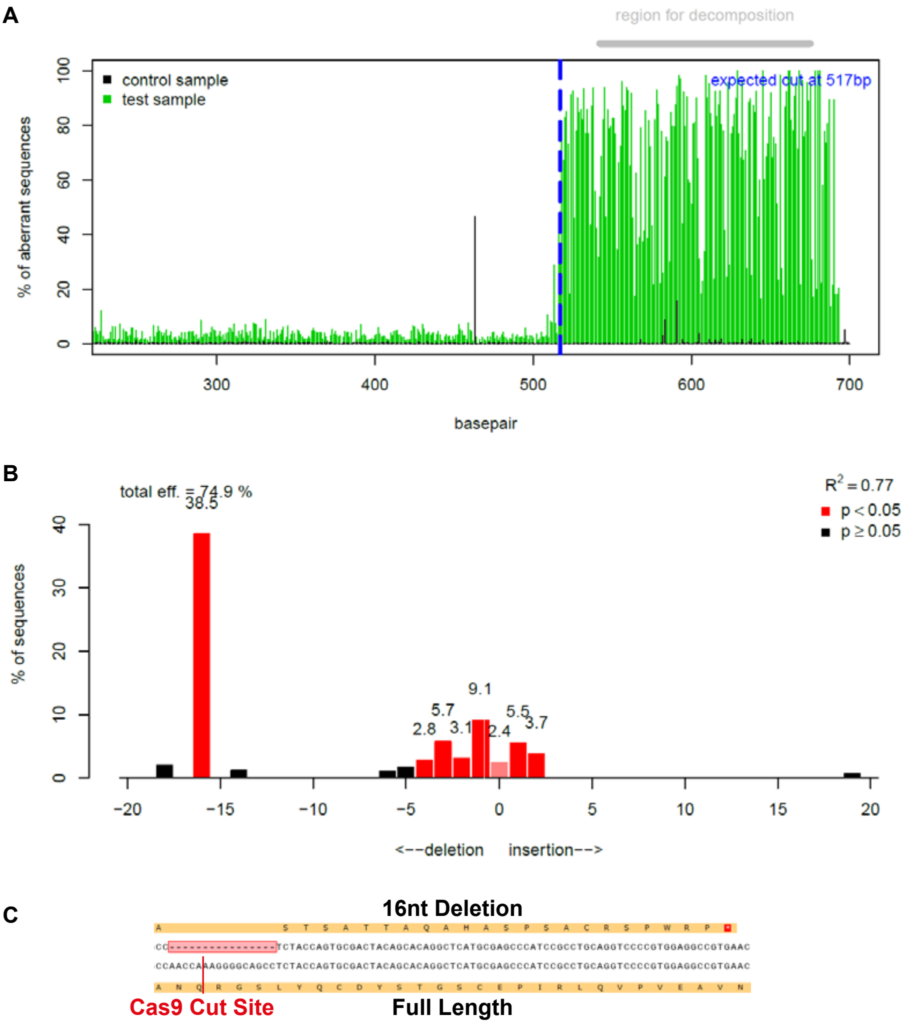

#### A.3 GapmeR Treatment of Differentiating CD34$^+$ cells

Antisense locked nucleic acid (LNA) GapmeRs (Qiagen) are single strand antisense DNA-RNA oligonucleotides which contain a central RNAse-H activating DNA moiety flanked by target-complementary sequences. These oligonucleotides cause RNAse-H mediated degradation of cytosolic target mRNA for potent gene silencing. We tested three separate GapmeR sequences targeting *ITGAM* and a negative control GapmeR for their ability to suppress CD11b in differentiating CD34$^+$ cells. We performed flow cytometric analysis during G-CSF directed differentiation (Fig. S3A) at a concentration of 3000 nM, which is the upper limit of the manufacturer's recommended working concentration. Media were changed every 48 hours and replenished with media containing G-CSF and fresh GapmeRs. CD11b positivity was assessed by flow cytometry using the gating shown in Fig. S1A. There was a nonspecific, low level suppression of CD11b expression during differentiation which recovered by day 14, where all GapmeR-treated cells expressed similar levels to untreated cells. qPCR analysis of *ITGAM* relative to undifferentiated CD34$^+$ cells did not show significant silencing at any chosen dose (Fig. S3B). Importantly, a positive control GapmeR targeting MALAT1 was ineffective at all concentrations, suggesting that GapmeR-mediated silencing is ineffective in differentiating CD34$^+$ cells (Fig. S3C). Testing the MALAT1 GapmeR in HeLa cells showed potent repression (data not shown) suggesting this lack of silencing was not due to the method or reagents used.

**Supplementary Figure 3**

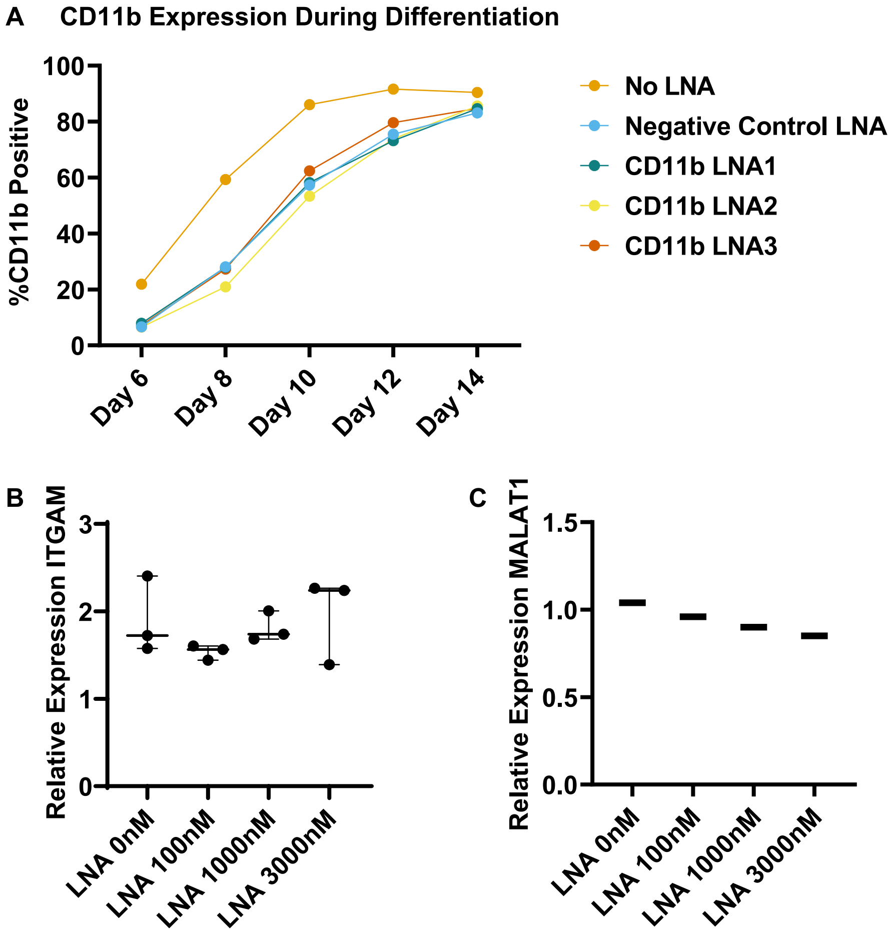

#### A.4 *ITGAM* guide RNA Selection

Fig. S4A shows histograms of CD11b expression gated per the strategy in Fig. S1D. Cells were expanded for 6 days in SFEM II and StemSpan supplement, electroporated with 0.4 nmol Cas9 complexed to 1 nmol of cr-tracr RNA targeting *ITGAM*, and then differentiated for 14 days in G-CSF. Guide sequence 1 was the most effective at suppressing CD11b expression in the differentiated $CD34^+$ cells. This guide was then tested in six distinct donors as shown in Fig S4B. and potently suppressed CD11b in four of the six donors tested.

**Supplementary Figure 4**

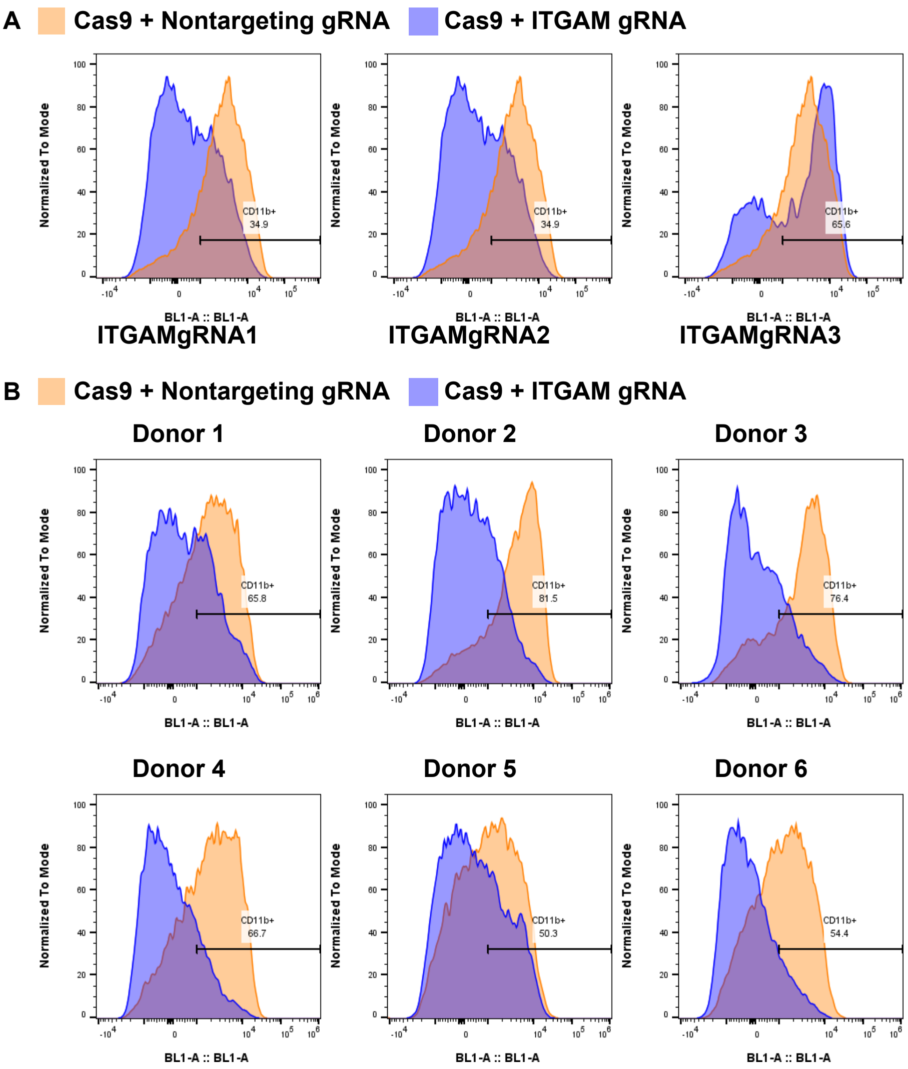

#### A.5 Gibson Assembly and Maps of Vectors

Fig. S5A shows the segmental map for pKLV-U6-shRNA-LacZ(BbsI)-PGKpuro2ABFP. Segment 1 was amplified from pLV-U6gRNA(BbsI)-PGKpuro2ABFP (Addgene #50946) using forward primer CTTGTGGAAAGGACGAAACACCGGGTCTTCTGTTGGCGGGTGTCGGGGC and reverse primer CCCTACCCGGTAGAATTGGATCCAGTCTTCAGCGAGTCAGTGAGCGAGGAAGC. Segment 2 was amplified from pUC19 using the forward primer ACAGAGGTTTACAAAATCAAGTGAGGTCGTTTCAACAAAAGGTAACCTGTTGGCGGGTGTCGGGGC and the reverse primer TGGTCGTTTGAGGGTAGAGAATTATTTAATGATGAGTATGGTGACCAGCGAGTCAGTGAGCGAGGAAGC. Fig. S5B shows the segmental map for pKLV-EFS-LacZ-PGK-Puro-tagBFP. Segment 1 was amplified from pLenti-spCas9-T2A-iRFP670-P2A-Puro (Addgene #122182) using forward primer GGACAGCAGAGATCCAGTTTGAAAGGAGTGGGAATTGGCTCC and reverse primer CTTTTGTTGAAACGACCTCACTTGATTTTGTAAACCTCTGTGTCCTGTGTTCTGGCGGCAAAC. Segment 2 was amplified from pUC.19 using forward primer ACAGAGGTTTACAAAATCAAGTGAGGTCGTTTCAACAAAAGCACCGGGTCTTCTGTTGGCGGGTGTCGGGGC and reverse primer TGGTCGTTTGAGGGTAGAGAATTATTTAATGATGAGTATGGATCCAGTCTTCAGCGAGTCAGTGAGCGAGGAAGC. Segment 3 was amplified from pLV-U6gRNA(BbsI)-PGKpuro2ABFP (Addgene #50946) using forward primer CATACTCATCATTAAATAATTCTCTACCCTCAAACGACCACAATTCTACCGGGTAGGGG and reverse primer GGAGCCAATTCCCACTCCTTTCAAACTGGATCTCTGCTGTCC. Fig. S5C shows the segmental map for pKLV-EFS-tagBFP-HincII-shRNA. This

plasmid was produced by using primers to excise the LacZ-PGK-Puro-2A segment from pKLV-EFS-LacZ-PGK-Puro-tagBFP using primers GTATAT-TATTCTAAGGTGTTCTTACCATGAGCGAGCTGATTAAGG (forward) and TCTACAGTTGTTAACTTGCCTTGTAAGTCATTGG (reverse) for segment 1 and ACAAGGCAAGTTAACAACTGTAGATCTTAGCCAC (forward) and GGTAAGAACACCTTAGAATAATATACGTCCTGT-GTTCTGGCGGC (reverse) for segment 2, for which the template in both instances is the original pKLV-EFS-LacZ-PGK-Puro-tagBFP plasmid. Fig. S5D shows the segmental map for pKLV-EFS-tagBFP-miR30a(BbsI). pKLV-EFS-tagBFP-HincII-shRNA was linearised at the HincII site to produce segment 1 using primers TGCCTACTGCCTCG-GACTTCAAGGGGGCTACTTTACTGTAGATCTTAGCCAC (forward) and CGCTCACTGTCAACAGCAATATACCTTCTTCTGCCTTGTAAGT-CATTGG (reverse). Segment 2 was amplified from pUC19 to incorporate a *LacZ*-containing golden gate flanked by miR-30a stem sequences into which miRshRNA oligonucleotides could be inserted using primers GAAGTCC-GAGGCAGTAGGCACCGTCTTCAGCGAGTCAGTGAGCGAGGAAGC (forward) and ATTGCTGTTGACAGTGAGCGACGTCTTCTGTTG-GCGGGTGTCGGGGC
(reverse).

**Supplementary Figure 5**

**A**

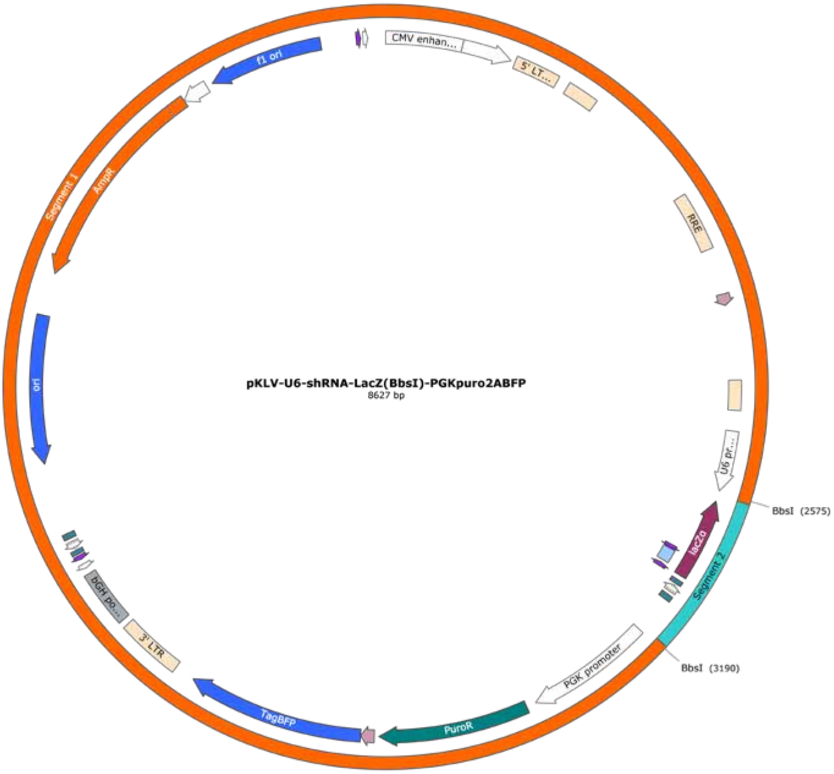

**Supplementary Figure 5**

**B**

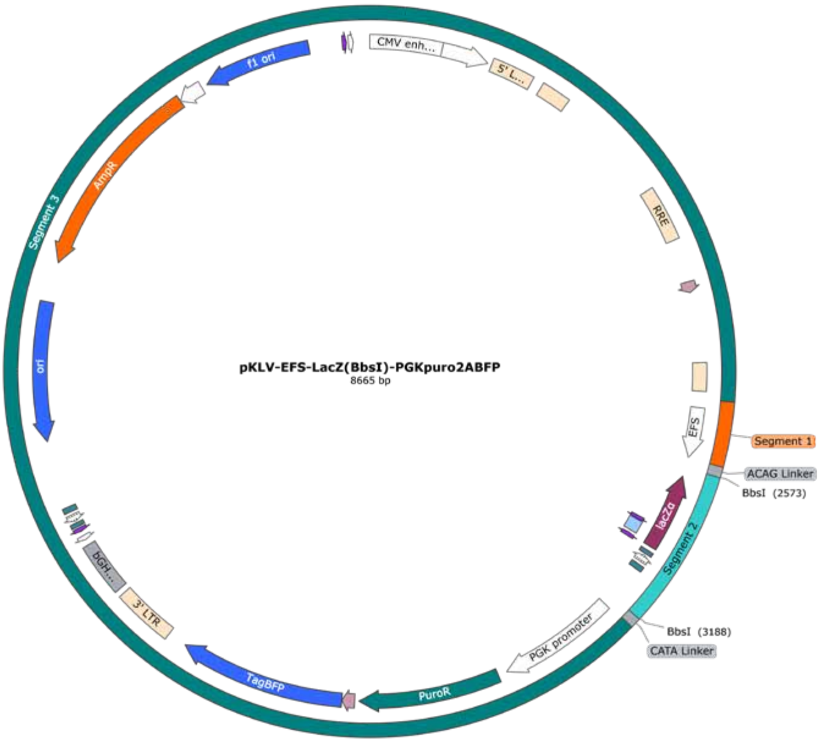

**Supplementary Figure 5**

**C**

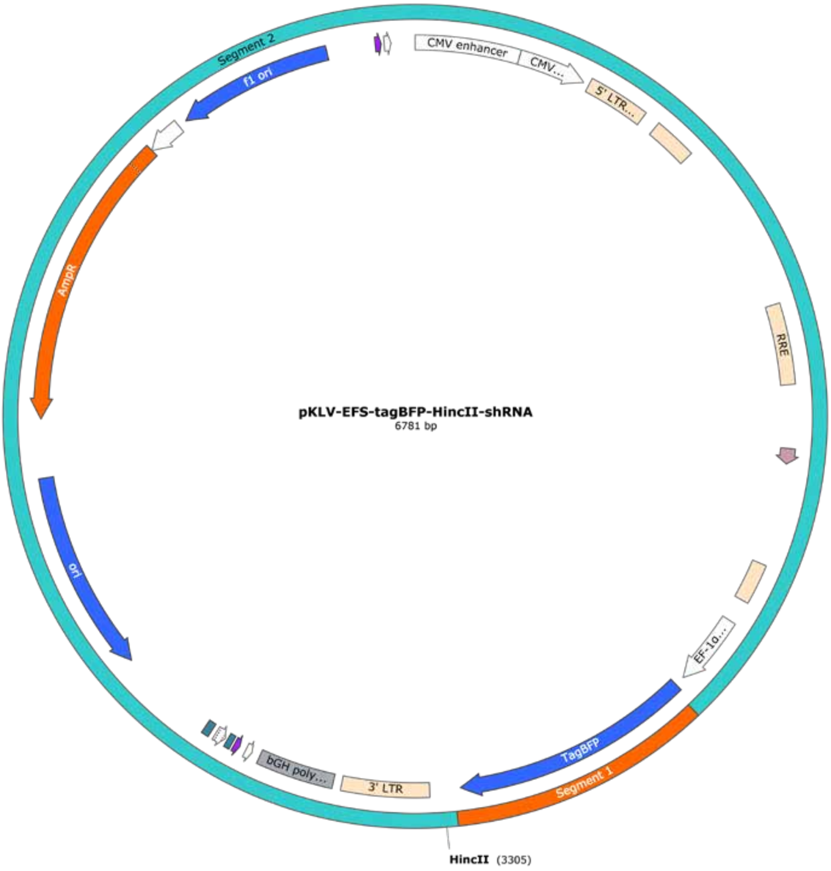

**Supplementary Figure 5**

**D**

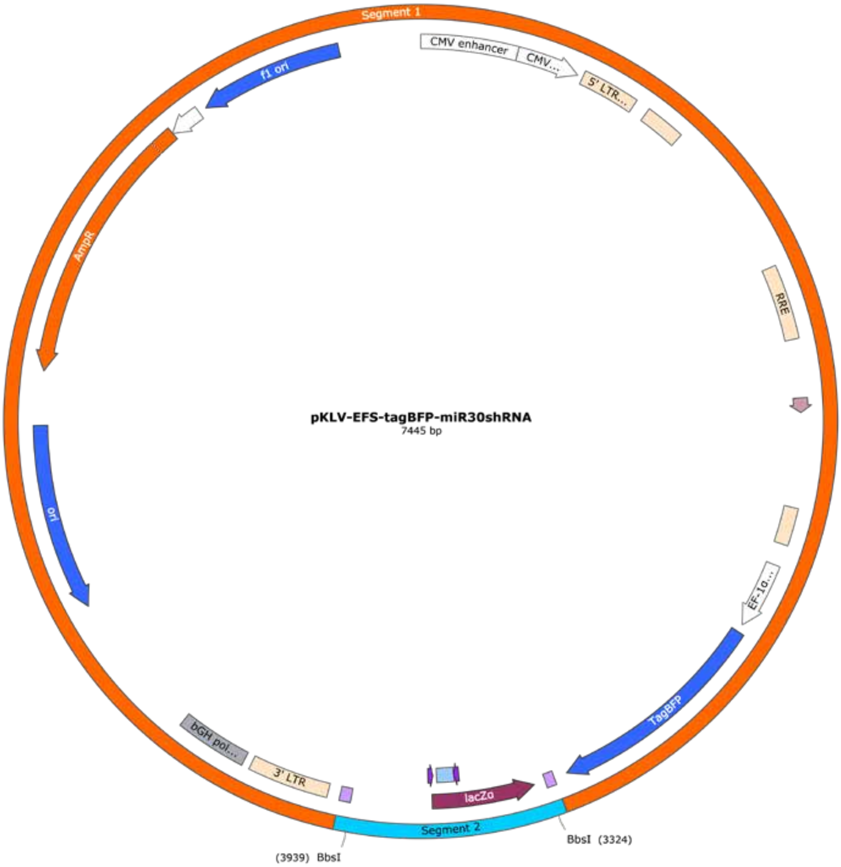

#### A.6 Plasmid List

pMD2.G (Addgene plasmid #12259; http://n2t.net/addgene:12259; RRID:Addgene_12259) and psPAX2 (Addgene plasmid #12260; http://n2t.net/addgene:12260; RRID:Addgene_12260) were both gifts from Didier Trono.

pLenti spCas9 T2A iRFP670 P2A puro was a gift from Raphael Gaudin (Addgene plasmid #122182; http://n2t.net/addgene:122182; RRID:Addgene_122182).

pKLV-U6gRNA(BbsI)-PGKpuro2ABFP was a gift from Kosuke Yusa (Addgene plasmid #50946; http://n2t.net/addgene:50946; RRID:Addgene_50946)[1].

pUC.19 plasmid DNA was obtained from New England Biolabs as part of commercially available reagent kits.

#### A.7 shRNA Sequences

| **Gene** | Type | **shRNA Name** | **shRNA Sequence** |
|---|---|---|---|
| Nontargeting | Pol III, miR21 loop | NTshRNA1 | GATCGACTGACAGATCGGAGCCCTGACCC-AGCTCCGATCTGTCAGTCGATCTTTTTT |
| ITGAM | Pol III, miR21 loop | ITGAMshRNA1 | GATATTAAGACATAAATTAGCCCTGACCC-AGCTAATTTATGTCTTAATATCTTTTTT |
| ITGAM | Pol III, miR21 loop | ITGAMshRNA2 | GTTAATAAATCAAATATAAGCCCTGACCC-AGCTTATATTTGATTTATTAACTTTTTT |
| ITGAM | Pol III, miR21 loop | ITGAMshRNA3 | GCAATGTGACTTTAATTTAGCCCTGACCC-AGCTAAATTAAAGTCACATTGCTTTTTT |
| Nontargeting | Pol II, miR30a Scaffold | NTsh1_miRE(30a) | GACGGGCGCAAAGGATATATAATAGTGAAGCCA-CAGATGTATTATATATCCTTTGCGCCCGTC |
| ITGAM | Pol II, miR30a Scaffold | ITGsh1_mirE(30a) | AGCGCCAGTCTTCTTTTGATATACTATAGTGAAGCCA-CAGATGTATAGTATATCAAAAGAAGACTGG |
| ITGAM | Pol II, miR30a Scaffold | ITGsh2_mirE(30a) | CTCAGACATCGGTTCATATTAATAGTGAAGCCA-CAGATGTATTAATATGAACCGATGTCTGAG |
| ITGAM | Pol II, miR30a Scaffold | ITGsh3_mirE(30a) | CTGGATTCATTTATTATTTCAATAGTGAAGCCA-CAGATGTATTGAAATAATAAATGAATCCAG |
| ITGAM | Pol II, miR30a Scaffold | ITGsh4_mirE(30a) | CTGAGTTAATAAATCAAATATATAGTGAAGCCA-CAGATGTATATATTTGATTTATTAACTCAG |
| ITGAM | Pol II, miR30a Scaffold | ITGsh5_mirE(30a) | ATCGGTTCATATTAAGACATAATAGTGAAGCCA-CAGATGTATTATGTCTTAATATGAACCGAT |

#### A.8 Guide RNA Sequences

| Gene | Guide RNA Name | Guide RNA Sequence |
|---|---|---|
| Nontargeting | NTgRNA1 | CATGCGCGCGCCATACCCTC |
| ITGAM | ITGAMgRNA1 | ATAGTGGCTGCCAACCAAAG |
| ITGAM | ITGAMgRNA2 | GGGCGGACACACACGGCCAC |
| ITGAM | ITGAMgRNA3 | CCGGTTCCGTAAGATGATGG |
